# Some plants don’t play games: An ideal free distribution explains the root production of plants that do not engage in a tragedy of the commons game

**DOI:** 10.1101/004820

**Authors:** Gordon G. McNickle, Joel S. Brown

**Affiliations:** University of Illinois at Chicago, Department of Biological Sciences, 845 W. Taylor St. (MC066), Chicago, IL, 60607, USA; Current address: Wilfrid Laurier University, Department of Biology, 75 University Avenue West, Waterloo, ON N2L 3C5, Canada

**Keywords:** tragedy of the commons, over-proliferation, ideal free distribution, competition, Evolutionary game theory, self/non-self recognition, Evolutionary stable strategies

## Abstract

This is the pre-peer-reviewed version of the following article: G.G. McNickle and J.S. Brown. (2014) An ideal free distribution explains the root production of plants that do not engage in a tragedy of the commons game. *Journal of Ecology.* DOI: 10.1111/1365-2745.12259, which has been published in final form at http://onlinelibrary.wiley.com/doi/10.1111/1365-2745.12259/abstract

1. Game theoretic models that seek to predict the most competitive strategy plants use for competition in soil are clear; they generally predict that over-proliferation of roots is the only evolutionarily stable strategy. However, empirical studies are equally clear that not all plants employ this strategy of over-proliferation of roots. Here, our goal was to develop and test an alternative non-game theoretic model that can be used to develop alternative hypotheses for plants that do not appear to play games.
2. The model is similar to previous models, but does not use a game theoretic optimization criterion. Instead, plants use only nutrient availability to select a root allocation strategy, ignoring neighbours. To test the model we compare root allocation and seed yield of plants grown either alone or with neighbours.
3. The model predicted plants that do not sense neighbours (or ignore neighbours) should allocate roots relative to resource availability following an ideal free distribution. This means that if a soil volume of quality *R* contains *x* roots, then a soil volume of quality *R*/*n* will contain *x*/*n* roots. The experimental data were consistent with this prediction. That is, plants grown with 1.2g of slow release fertilizer resources produced 0.043 g of roots, while plants grown with neighbours, or plants grown with half as much fertilizer produced half as much root mass (0.026g, and 0.24g respectively). Seed yield followed a similar pattern.
4. This model presents an alternative predictive framework for those plant species that do not seem to play a tragedy of the commons game for belowground competition.
5. *Synthesis:* It remains unclear why some plants do not engage in belowground games for competition. Models suggest over-proliferation is an unbeatable evolutionary stable strategy, yet plants that do not play the game apparently coexist with plants that do. We suggest that a greater understanding of trade-offs among traits that are important for other biotic interactions (above-ground competition, enemy defence, mutualisms) will lead to a greater understanding of why some species over-proliferate roots when in competition but other species do not.

## INTRODUCTION

Belowground competition among plants can be both important and intense (Casper and Jackson 1997; Schenk 2006). In many terrestrial systems, plants are strongly limited by soil nitrogen availability (Robinson 1994; Vitousek and Howarth 1991) and competition belowground can reduce plant growth and performance by several orders of magnitude (Lamb and Cahill 2008; Wilson 1988). As a result competitive interactions among plants in soil can have important consequences for the ecology (Cahill and McNickle 2011; Casper and Jackson 1997) and evolution (Gersani et al. 2001; Thorpe et al. 2011) of plants. Thus, it is logical to ask: what rooting strategies lead to the greatest competitive ability and highest returns on fitness? Cahill and McNickle 2011; Herben and Novoplansky 2010; Litav and Harper 1967; Parrish and Bazzaz 1976; Schenk et al. 1999; Litav and Harper 1967; Craine et al. 2005; Gersani et al. 2001, Campbell and Grime 1989; Parrish and Bazzaz 1976; Parrish and Bazzaz 1976; Schenk et al. 1999; Hess and de Kroon 2007; Litav and Harper 1967; McNickle et al. 2008; Mommer et al. 2011; Schenk et al. 1999.

Casper and Jackson 1997; Goldberg et al. 1999; Schenk 2006 Casper and Jackson 1997; Schenk 2006; Tilman 1982Wilson 1988 McNickle et al. 2008; Mommer et al. 2011; Schenk 2006Jackson et al. 1996; Jackson et al. 1997Jackson et al. 1997Jackson et al. 1997Tilman 1982; Tilman et al. 1997Dudley and File 2007; Falik et al. 2003; Gruntman and Novoplansky 2004; Mommer et al. 2010; Padilla et al. 2013 Cahill and McNickle 2011; Cahill et al. 2010Jackson et al. 1996; Jackson et al. 1997Cahill and McNickle 2011; Cahill et al. 2010; Mommer et al. 2010; Parrish and Bazzaz 1976; Schenk et al. 1999; von Felten and Schmid 2008Parrish and Bazzaz 1976Plant ecologists seem to be divided on this question. In one group, game theoretic models of root competition suggest that all else equal, plants should exhibit plasticity and over-proliferate roots in the presence of a neighbour relative to when grown alone (Craine 2006; Craine et al. 2005; Dybzinski et al. 2011; Gersani et al. 2001; McNickle and Brown 2012; McNickle and Dybzinski 2013; O’Brien and Brown 2008; O’Brien et al. 2007).McNickle and Brown 2012 The prediction of over-proliferation seems to be an extremely robust prediction even with a wide variety of different model formulations, as long as a game theoretic optimization criterion is used. In these game theoretic models plants must possess two abilities: (i) the ability to sense and respond to nutrient availability (e.g. Forde and Walch-Liu 2009; Ho et al. 2009; Zhang and Forde 1998; Zhang et al. 1999) and (ii) the ability to sense and respond neighbour stratgies (e.g. Falik et al. 2003; Falik et al. 2005; Gersani et al. 1998; Gruntman and Novoplansky 2004). There is good evidence that at least some plants possess both abilitiesDybzinski et al. 2011. Thus, when plants can sense both neighbours and nutrients their best strategy is to produce more roots than are necessary to harvest the available nutrients in an attempt to pre-empt the resource supply of neighbours. Theory is clear that this is an evolutionarily stable strategy because it cannot be invaded by a less proliferative strategy.

A second group of plant ecologists are skeptical of these game theoretic models of root production (de Kroon et al. 2012; Herben and Novoplansky 2010; Hess and de Kroon 2007; Laird and Aarssen 2005; Schenk 2006; Semchenko et al. 2007). This skepticism seems to be well founded, and there is evidence that many plants do not engage in these plastic pre-emptive games (Cahill et al. 2010; Semchenko et al. 2007; Semchenko et al. 2010). In general the criticisms seem to amount to questioning whether plants possess the ability to sense neighbours (Hess and de Kroon 2007). If plants cannot sense neighbours then it follows that game theoretic models of plant plasticity in root production do not apply because it is impossible for such plants to respond to neighbours and play these behavioural games. This skepticism might be well founded, as currently the mechanisms that might allow plants to sense neighbours remain a mystery (McNickle and Brown 2012; Novoplansky 2009). Given these two alternative theoretical expectations, what patterns do we observe in data?

Empirical tests of these game theoretic models are clear: plant species are mixed in their responses to nutrients and neighbours. There is good evidence that some plants play root games that lead to over-proliferation of roots (de Kroon et al. 2012; Gersani et al. 2001; Gruntman and Novoplansky 2004; Lang’at et al. 2013; Maina et al. 2002; Mommer et al. 2010; O’Brien et al. 2005; Padilla et al. 2013, note that some authors have begun using the term over-yielding instead of over-proliferation), and there is equally good evidence that some plants do not play these games (Cahill and McNickle 2011; Cahill et al. 2010; Litav and Harper 1967; Schenk et al. 1999; Semchenko et al. 2007; Semchenko et al. 2010). In the “game-on” camp, a variety of game theoretic models have been proposed and analyzed (Craine 2006; Craine et al. 2005; Dybzinski et al. 2011; Farrior et al. 2013; Gersani et al. 2001; McNickle and Brown 2012; O’Brien and Brown 2008; O’Brien et al. 2007). However, despite the large amount of criticism of these game theoretic models (de Kroon et al. 2012; Hess and de Kroon 2007; Laird and Aarssen 2005; Schenk 2006; Semchenko et al. 2007), the “game-off” camp has not produced many alternative models to predict the root production strategy of plants that do not engage in a tragedy of the commons game. Given that many plants do not seem to engage in these root games, there would be value in asking: what is the best root allocation strategy for a plant that cannot sense neighbours and as a result cannot play pre-emptive games?

In this paper, we propose an alternative model for plants that do not engage in a tragedy of the commons game. We test this with fast cycling *Brassica rapa* (L., var Wisconsin Fast Plants^®^, Carolina Biological Supply Company, Burlington, NC, USA). We asked; (1) what is the best strategy for root production of a plant that can sense nutrient availability but cannot sense neighbours and faces competition? To address this question, we develop a simple model that is similar to previous game theoretic models (McNickle and Brown 2012; O’Brien et al. 2007), but uses a non-game theoretic optimization criterion. (2) How does *B. rapa* allocate to roots when grown alone or when grown with neighbours and how do these root allocation strategies influence reproductive output? Based on our experience with this plant (GGM Personal Observation), we did not expect this plant to be capable of sensing neighbours and therefore it should not be capable of engaging in a tragedy of the commons game. (3) How does the set of best responses of *B. rapa* compare to either a “game-on” prediction or a “game-off” prediction of focal plant root production strategies relative to neighbour root production strategies. Given that the world appears to contain a mixture of game on and game off plants, we discuss some implications of this at the end of the manuscript.

## METHODS

### Model

Here we develop a model of root allocation for plants that respond to nutrients only. These plants can still experience competitive effects due to depletion of the resource environment by neighbours, but in this model plants cannot sense neighbours and so they do not take into account the competitive strategies of their neighbours when allocating to root biomass. That is, they employ a resource matching strategy and do not play a game. Instead, as neighbours depress the nutrient environment these “game off” plants simply perceive the environment as becoming worse, and produce fewer roots in the presence of a neighbour compared to alone. This model is similar to our previously published game theoretic models (Gersani et al. 2001; McNickle and Brown 2012; O’Brien and Brown 2008; O’Brien et al. 2007), but the model presented here is not game theoretic.

Let *R* be the available resources in the soil, let *u_f_* be the root production of the focal plant*, u_n_* be the root production of the neighbour plant and let *r* be the sum total amount of roots of all competing plants (*r* = *u_f_* + *u_n_*). Further, let, *c* be the per-unit cost of root production for all plants. We assume diminishing returns of nutrient harvest with increasing root production. We also assume that root production costs and each plant’s share of the nutrient harvest increase monotonically with total root production. Finally, we assume that there is no shoot competition and that plants are not light limited. Thus, the net harvest (π_*f*_) of an annual plant at senescence can be represented as, their share of the nutrient harvest minus the costs of root production, or,

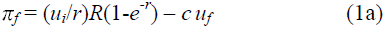

We write this for the focal plant only, but an identical equation exists for the neighbour plant with the subscripts reversed. Plants that can sense neighbours and play a plastic tragedy of the commons game (McNickle and Brown 2012) will choose a root production strategy, 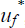, that satisfies,

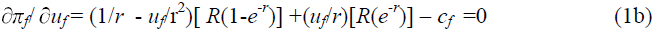

We present the game theoretic version of this model here purely as a point of comparison with the non-game theoretic version. However, we do not analyze it in detail; it is described and analyzed in detail in McNickle and Brown (2012). For plants that cannot sense neighbours and do not play a plastic tragedy of the commons game we need an alternative optimization criterion which is not game theoretic (Anten and During 2011). We hypothesise that these “game-off” plants should be blind to the strategy of neighbours (or at least ignore it) and as a result these plants should not base their root production strategy on their expected share of the nutrients. As a result, we propose that such game off plants should ignore the term *u_f_*/*r* in equation 1a, and focus only on the resource environment, any depletion trajectories (caused by self or non-self) and their own costs associated with root production. Thus, we define an expected payoff, *P_f_*, equation based on eqn 1a that omits this term,

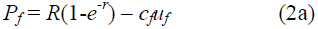

This is not the plant’s actual payoff, but it is the payoff that a plant who is blind to the presence of neighbours will expect since it can only detect the resource environment but not the presence of neighbours. Note that these “game-off” plants can still sense any resource depletion that is caused by neighbours, however they are incapable of distinguishing between resource depletion caused by their own roots or the roots of neighbours. Thus, the optimal root production strategy, 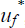, of a plant that cannot detect neighbours should satisfy,

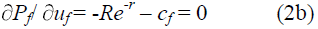

Solving equation (2b) for *u_f_\** gives the optimal root production strategy of plant *f,* and a version of equation (2b) must be simultaneously solved for plant *n*. However, though these game-off plants expect the payoff given in equation (2a), and even though they don’t sense the neighbour, they still must compete with neighbours and their actual payoff is still given by equation (1a) and not by equation (2a).

### Experiment: What is the proper control?

To test the model we grew *B. rapa* plants either alone or with neighbours. Plants were grown in three treatments based on model expectations (Equations 1-2), past empirical tests of game theoretic models (e.g. Gersani et al. 2001; Maina et al. 2002; O’Brien et al. 2005) and criticisms of those experiments (Hess and de Kroon 2007; Laird and Aarssen 2005; Schenk 2006; Semchenko et al. 2007). Early experiments testing for a tragedy of the commons game were interested in root allocation based upon resource availability. Thus, these experiments attempted to control resource supply per-plant (*R*) but they did so by manipulating pot volume (*V*). For example, plants with neighbours were grown in pots of volume *V* containing *R* resources, and plants alone in pots of volume *V*/2 containing *R*/2 resources (e.g. Gersani et al. 2001; O’Brien et al. 2005). This design was based on the hypothesis that plants which do not respond to neighbours should respond only to nutrient availability (e.g. equation 1b vs 2b) and thus it was critical to control nutrient availability per-plant to test whether plants respond only to nutrient availability or to both nutrient availability and neighbours (Gersani et al. 2001; McNickle and Brown 2012). However, though this design controls resource availability at the per-plant level, it does not control pot volume.

In pot experiments, volume can affect plant growth if plants become pot-bound during the course of the experiment. Thus, many authors have been critical of this method for controlling nutrient availability and argue that it is more important to control pot volume than nutrient availability (Hess and de Kroon 2007; Schenk 2006). However, controlling only pot volume produces a design where plants grown alone have access to pots of volume *V* and *R* nutrient resources, while plants grown with neighbours have access to pots of volume *V* but only *R*/2 resources. In this design plants grown alone are almost always significantly larger than plants grown with neighbours because they have been given access to twice as many nutrients making it difficult to compare plants grown alone to plants grown with neighbours. The appropriate control for these types of competition experiments comes down to whether one expects pot volume to be a more important determinant of plant growth and competition, or whether one expects nutrient availability to be a more important determinant of plant growth and competition. In reality, the best approach is probably to employ both controls.

Here, we used both a volume and a nutrient availability control. Plants were grown with neighbours in pots of volume *V* (*V*=1L, 10×10×10cm), and plants grown alone were either grown in pots of volume *V* or *V*/2 (Figure 1). This let us compare the ‘with neighbours’ treatment to both controls for either volume or nutrient availability per plant.

**FIGURE 1:**
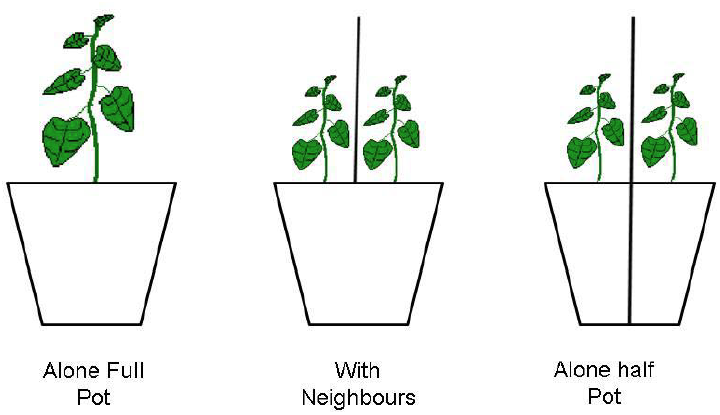
Schematic of experimental design. Plants were either grown in three treatments; (*i*) alone in full sized pots of volume *V* = 1L, (*ii*) with neighbours in pots of volume *V*, or alone in pots of volume *V*/2. Opaque screens were erected between paired plants so that they did not compete or interact aboveground.

### Growing conditions

Soil was a 1:3 mixture of potting soil (Miracle-Gro^®^ Moisture Control^®^ Potting Mix, The Scotts Company LLC, Marysville, OH, USA) to washed sand (Quickrete, Atlanta, GA, USA) which were each pre-seived through a 2mm screen to facilitate root extraction at harvest. Soils were amended with 19-6-12 (N-P-K) slow release fertilizer pellets on the surface of the soil (Osmocote, The Scotts Company LLC, Marysville, OH, USA). Fertilizer was supplied based on an equal supply per volume of soil, such that full sized pots (alone full pot, and with neighbours treatments) each were given 1.2 grams of fertilizer while half sized pots were given 0.6 grams. Plants were grown in a controlled growth chamber with continuous fluorescent lighting. Plants were spaced at least 15 cm apart on the bench, and opaque screens were erected between competing plant so they did not interact aboveground in any treatments. This forced all competitive interactions to occur belowground, in order to satisfy the assumption of the model that there was no shoot competition. Additionally, plants in each treatment were grown in pairs (with neighbours or alone) where each plant was *a priori* designated as either focal or neighbour which produced pairs of plants that experienced the same conditions on the bench. Plants were watered daily using an automatic drip irrigation system that supplied ∼126mL of water per day to pots of volume *V*, and ∼63mL of water per day to pots of volume *V*/2. Because we were trying to generate best response curves of each plants (see below), each alone treatment included 25 pairs of plants, and the ‘with neighbours’ treatment included 50 pairs of plants (total of 100 focal plants, and 100 neighbour plants).

On day zero, multiple seeds were sewn per planting location, and these were thinned to one plant per location within 1 day of germination. Plants were grown for 40 days under continuous lighting conditions which is recommended for fast cycling *B. rapa* (Carolina Biological Supply Company, Burlington, NC, USA). Location on the bench was re-randomized every 5 days to minimize bench effects. *B. rapa* is self-incompatible, and fruit takes approximately 20 days to mature. All flowers produced up to day 20 were hand pollinated using cotton swabs. After 20 days pollination was discontinued and fruit were permitted to mature for 20 more days. Mates were haphazardly chosen each day, so each plant had a new mating partner on each day of pollination. After 40 days of growth plants had begun senescence and fruits, leaves and stems were harvested. Roots were washed on a 2mm sieve, and the root systems of neighbouring plants were carefully separated by floating them in water and gently shaking them to separate. We had no difficulty carefully separating the roots of each plant grown in competition by marking the stem of each plant, and gently shaking the root system in water until they separated. Subsequently, biomass was dried at 60°C to constant mass, and weighed. Fruits were dried at room temperature, counted and weighed. Fruits were then opened and seeds were counted and weighed. All statistical models were run in the R statistical environment (R-Development-Core-Team 2009).

## RESULTS

### Model results, and best response curves

McNickle and Brown (2012) introduced the concept of a best response curve that plots one plant’s root production strategy, against the root production strategy of its neighbour and we use this as one diagnostic criterion to test whether plants are playing a game. From our model (Equation 1, 2) this would plot the points 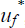 vs ***u_n_*** for the focal plant’s best response to any possible strategy of the neighbour, and ***u_f_*** vs *u_n_\** where **u_n_** and **u_f_** are continuous vectors of all possible strategies either plant might choose. This set of best responses can be thought of as a competitive “play book” for each plant: if the neighbour produces *y* roots, the best response of the focal plant is to produce *x* roots and *vice versa*. Comparing the game theoretic, and non-game theoretic version of this nutrient competition model, the best response curves from each model are quite different and this can be a useful diagnostic tool for experimentally comparing game on or game off plants (Figure 2).

**Figure 2:**
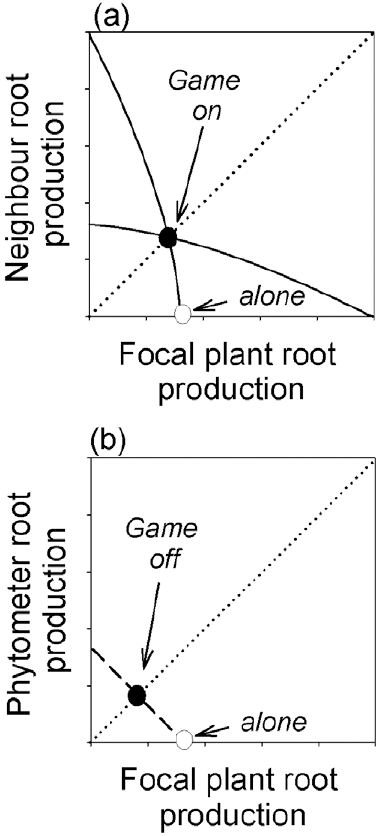
Example best response curves for game on plants (a) or game off plants (b). For plants that engage in a tragedy of the commons game, they over-proliferate roots in response to neighbours causing the best response curve to bow up and out from the origin (panel a). Game on plants will produce more roots in the presence of neighbours compared to alone (potentially much more than we’ve shown). For plants that do not engage in a tragedy of the commons game the model predicts a linear best response curve that obeys an ideal free distribution of roots relative to soil nutrient concentrations (b). Game off plants will never produce more roots when neighbours are present compared to alone.

Based on the model, the root production strategy of the non-game theoretic model presented here follows an ideal free distribution (Fretwell and Lucas 1969, Gersani et al. 1998). This means that if a soil volume of quality *R* contains *x* roots, then a soil volume of quality *R*/*n* will contain *x*/*n* roots. Similarly, if a plant grown alone in a soil volume of quality *R* produces *x* roots, then a plant grown with *N* neighbours in a soil volume of quality *R* should produce *x*/*N* roots. The best response curve of intraspecific competition among game-off annual plants is shown in Fig 1b. It is a straight line connecting the root production strategy of each plant when grown alone. Applied to our experimental treatments, the model predicts that each plant should produce exactly half as many roots when grown with neighbours in pots of volume *V* as when grown alone in pots of volume *V*. Additionally, plants grown with neighbours in pots of volume *V* should each produce exactly the same amount of roots as plants grown alone in pots of volume *V*/2.

Alternatively, game-on plants choose a root production strategy based on both the resource environment, and the strategy used by neighbours (equation 1b). This causes game-on plants to over-proliferate roots in an attempt to pre-empt the resource supply of neighbours, and several versions of these game theoretic models have been described and analyzed in detail elsewhere (Craine et al. 2005; Dybzinski et al. 2011; Gersani et al. 2001; O’Brien and Brown 2008; O’Brien et al. 2007) including an analysis and discussion of best response curves (McNickle and Brown 2012). An example best response curve of intraspecific competition among game-on annual plants is shown in Fig 1c (McNickle and Brown 2012). Here, curved lines bow-out from the origin because of the strategy of over-proliferation causes plants to produce significantly more roots in the presence of neighbours compared to when they grow alone (McNickle and Brown 2012). In the context of our experiment, a game theoretic model would predict that, plants grown with one neighbour in pots of volume *V* should produce significantly more than half the roots of a plant grown alone in a pot of volume *V*, and significantly more roots than plants grown alone in pots of volume *V*/2.

### Experimental results

In this study, *B. rapa* did not engage in a tragedy of the commons game (Figure 3). One way ANOVA revealed that plants grown with neighbours had exactly the same seed yield and root biomass as plants grown in half volume pots, and exactly half of the seed yield (F_2,147_=29.69, p<0.0001) and root biomass (F_2,147_=49.01, p<0.0001) as plants grown in full volume pots (Figure 3a, b). Patterns of leaf, stem and fruit production followed the exact same pattern as Figure 3a and Figure 3b, and are not shown. When plotted as best responses, the observed root production strategies of *B. rapa* fall on the ideal free distribution line which reject the game theoretic model and support the simpler game-off model for this species (Figure 3c,d).

**Figure 3:**
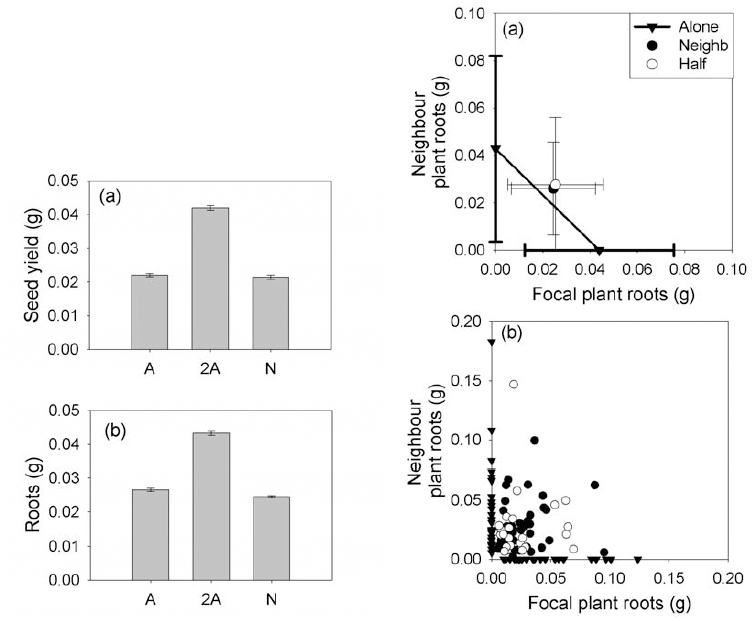
Mean seed yield (a) and root mass (b) of focal plants grown either alone in pots of volume V/2 (A), alone in pots of volume V/2 (A/2) or with neighbours (N). These data are also plotted as best response curves (c-d). Mean responses of plants grown with neighbours or alone in half pots are compared to the expected best response curve (a). Raw data are shown in panel (b). Error bars are ± 1 SD.

## DISCUSSION

Simple game theoretic models that include only root growth and hold all else equal predict that a plants best response is to always over-proliferate roots in the presence of a neighbour (Gersani et al. 2001; McNickle and Brown 2012). In these models over-proliferation of roots is an evolutionary stabile strategy in that it should not be invasible by plants that use a less proliferative strategy. The empirical evidence show that some plant species clearly do play a tragedy of the commons game when choosing a root production strategy (Cahill and McNickle 2011; Falik et al. 2003; Mommer et al. 2010; Padilla et al. 2013; Semchenko et al. 2010). However, evidence also shows that other species clearly do not play such games (Figure 3, Cahill et al. 2010; Schenk et al. 1999; Semchenko et al. 2010). The invasability criterion of an ESS would imply that these two strategies should not be capable of coexisting in natural systems. Yet again, the available data seem to suggest that these plants may coexist within the same system (Cahill and McNickle 2011; de Kroon et al. 2012; Mommer et al. 2010; Semchenko et al. 2010). This suggests that there is a gap in our theoretical understanding of plant root production strategies.

One way of thinking of this gap is that some species (game off plants) may simply not possess the ability to sense neighbours, while other species (game-on plants) do possess this ability. This is the proximate cause of the lack of any game theoretic response, that is, plants simply lack the proximate mechanisms to permit them to play such games. However, this leaves the ultimate evolutionary cause of this lack of game theoretic responses unanswered: why has evolution apparently furnished some species with the ability to sense neighbours, but failed to furnish other species with this ability. This is an especially important question to ask since a wide variety of models differing greatly in their parameterization, complexity and formulation have all consistently shown that the game-theoretic response of over-proliferation of roots is the only evolutionarily stable strategy for root competition (Craine 2006; Craine et al. 2005; Dybzinski et al. 2011; Gersani et al. 2001; McNickle and Brown 2012; McNickle and Dybzinski 2013; O’Brien and Brown 2008; O’Brien et al. 2007).

Most current game theoretic models employ the simplifying assumption that competition belowground is the only important process for plant fitness and this assumption is only met in the most controlled of glasshouse experiments (E.g. Figure 1). Yet, the evolutionary history of plants requires trade-offs between traits associated with root and shoot competition, investment into mutualisms, defence against enemy attack and a myriad of biophysical and environmental pressures. All of these biotic interactions become a tragedy of the commons when modeled using evolutionary game theory (McNickle and Dybzinski 2013), yet most models focus only on one process, holding all others equal, and predict a single ESS solution. At present we lack a good understanding of how plants make trade-offs between investments in each biotic interaction, and how investment in one biotic interaction shifts investment into other biotic interactions. For example, when should a plant invest in enemy defence instead of belowground competitive ability? When should a plant invest in mutualisms instead of enemy defence? We currently lack good answers to these questions (McNickle and Dybzinski 2013). However, one hypothesis is that those plants which do not engage in a tragedy of the commons game for root competition, might be adapted to focus more strongly on other interactions which are important in plant ecology. This could include aboveground competitive ability, defence from enemy attack or investment into mutualistic associations (Archetti et al. 2011; Falster and Westoby 2003; McNickle and Dybzinski 2013; Oksanen 1990). We suggest that understanding why some plant species are adapted to be capable of sensing neighbour roots and engaging in a tragedy of the commons game while other plant species clearly cannot sense neighbour roots and cannot engage in such games, will be an important question to move this debate on what strategies maximize plant competitive ability belowground forward.

The idea that root growth of plants might follow an ideal free distribution, is not new (Gersani et al. 1998). Gersani et al. (1998) presented a graphical model based on Fretwell’s (1972) fitness density model that used nutrient uptake per unit root to estimate root allocation of plants via density dependent. They tested this graphical model using *Pisum sativum*, L. and showed that plants with roots evenly split between one pot containing five competitors and a second neighbour-free pot would preferentially allocate roots into the empty pot away from neighbours. Moreover, these peas followed an ideal free distribution in terms of their root allocation (i.e. *x* roots in the alone pot, and *x*/2, *x*/3, or *x*/5 roots in pots with 2, 3 or 5 neighbours respectively). Interestingly, *P. sativum* seems to be among those plants which are capable of sensing neighbours and engages in a strategy of over-proliferation of roots in the presence of neighbour plants (Falik et al. 2003). This suggests that not only is there a mixture of “game-on” and “game-off” strategies among plant species, but that also within a species that is capable of detecting neighbours, plants may sometimes use a root allocation strategy of over-proliferation leading to a tragedy of the commons game and sometimes use a more restrained strategy leading to an ideal free distribution of roots. Why and how plants might switch strategies remains unknown, and will be an interesting question for plant ecologists interested in strategies for root competition.

The hypothesis of an ideal free distribution also appears to be a common null hypothesis, though it is not typically named as such. For example, Padilla et al (2013) grew *Festuca rubra* and *Plantago lanceolata* either in monocultures or in mixtures. Their null hypothesis was that the root production of each plant when grown in mixture would be exactly ½ of the observed root production of each monoculture. This amounts to an expectation that roots follow an ideal free distribution (Figure 2, Gersani et al. 1998). Padilla et al (2013) rejected this null hypothesis and found that plants in mixture produced significantly more roots than were expected. They called this over-yielding, but in the context of previous game theoretic ideas it could be called over-proliferation of roots (Gersani et al. 2001). Mommer et al (2010) performed a similar experiment using monocultures and mixtures using the same null hypothesis, and also found over-proliferation of roots in mixture compared to the null expectation of an ideal free distribution (they did not call it an ideal free distribution, and also used the term “over-yielding” for reasons that are unclear to us).

If these ideas are already implicit in the plant competition literature, what do we gain by explicitly recognizing that root growth of “game-off” plants might follow an ideal free distribution? We suggest that explicitly modeling root allocation of plants that do not respond to neighbours as an ideal free distribution adds formalism and predictive power to theories of competition. The ideal free distribution has well understood properties (Fretwell and Lucas 1969), and many ideas in ecology are built around this distribution (Křivan et al. 2008; Sutherland 1983) which might shed light on plant strategies for competition. The model presented here provides a clear alternative model to the tragedy of the commons models of root production (e.g. Gersani et al. 2001) which have been so sharply criticised by so many authors (E.g. Hess and de Kroon 2007; Schenk 2006). An alternative model can focus thinking and experiment in a way that the ongoing discussions have not. Though for those plant ecologists that have argued against game theoretic applications to problems in plant ecology, it is worth recognizing that behaviours which generate an ideal free distribution of organisms among habitats can, themselves, be an evolutionarily stable strategy for dispersal derived from a game theoretic model (Křivan et al. 2008).

Finally, when thinking about a tragedy of the commons evolutionary game, one should also consider the distinction between a fixed allocation to roots and plastic allocation to roots. The game presented here is one of plasticity, it can be thought of as a behavioural game where plants who can sense neighbours (Falik et al. 2003; Gruntman and Novoplansky 2004) should respond differently to neighbours compared to plants that cannot sense neighbours (Gersani et al. 2001; McNickle and Brown 2012). However, whether plants play a behavioural game via plasticity in allocation during the lifetime of a single plant, is a different question from whether they play a fixed allocation game in evolutionary time (McNickle and Dybzinski 2013). Evolutionary time leads to an evolutionary arms race for allocation to resource harvesting organs and organs associated with competitive ability. For example, such an arms race has produced extremely tall woody plants that have a high fixed allocation to wood over evolutionary time due to tragedy of the commons games for light competition (Falster and Westoby 2003; Givnish 1982; Givnish 1995; King 1990; Oksanen 1990). Though less well recognized, a similar evolutionary arms race has likely influenced fixed allocation to roots by plants in evolutionary time (Dybzinski et al. 2011; McNickle and Dybzinski 2013). That is, modern plants (including *B. rapa*) may have a fixed allocation to root biomass that is higher than their ancestors due to an evolutionary arms race, even while they cannot sense neighbours and do not engage in plastic behavioural games during the course of their lifetime. This fixed allocation game would produce “game-off” best response curves exactly as shown in Figure 1b over the course of one plant’s life. However, over evolutionary time the expectation would be that plants would gradually increase total root allocation, causing the linear best response curve to move further and further from the origin, even while maintaining an ideal free distribution within each generation. Unfortunately, testing this fixed allocation arms race game will be significantly more challenging and can likely only be achieved with labour intensive artificial selection experiments.

## Conclusions

Evolutionary game theory has injected some controversy into our understanding of the best strategies plants may use for competition underground. Currently, the data are quite mixed and suggest that some plants play behavioural games with their roots, and others do not. Even within a plant species some contexts seem to lead to behavioural games while others do not. Here we presented and tested an alternative non-game theoretic model of root allocation strategies for plants that compete for nutrients but cannot sense neighbours (or at least ignore neighbours). The model predicts that in this case, root production should follow an ideal free distribution. The model plant *B. rapa* which does not appear to sense neighbours responded as predicted by this “game-off” model, reducing root production relative to resource availability regardless of whether this was caused by pot size, or depletion by neighbours. That root allocation might follow an ideal free distribution is not a new idea, but our model adds formalism to some existing ideas in the competition literature. Further, our model provides a clear alternative model for those species which do not appear to engage in a tragedy of the commons game.

## ACKNOWLEDGEMENTS

GGM thanks the Natural Sciences and Engineering Research Council of Canada for a Post-Doctoral Fellowship and for a Banting Post-Doctoral Fellowship. We thank MA Gonzalez-Meler for access to growth facilities, CJ Whelan and P Orlando for discussions on experimental design and interpretation, and P. Lim and R.J. Kilgour for assistance with experimental setup and harvest.

